# Isoform and pathway-specific regulation of post-transcriptional RNA processing in human cells

**DOI:** 10.1101/2024.06.12.598705

**Authors:** Karan Bedi, Brian Magnuson, Ishwarya Venkata Narayanan, Ariel McShane, Mario Ashaka, Michelle T. Paulsen, Thomas E. Wilson, Mats Ljungman

## Abstract

Steady-state levels of RNA transcripts are controlled by their rates of synthesis and degradation. Here we used nascent RNA Bru-seq and BruChase-seq to profile RNA dynamics across 16 human cell lines as part of ENCODE4 Deeply Profiled Cell Lines collection. We show that RNA turnover dynamics differ widely between transcripts of different genes and between different classes of RNA. Gene set enrichment analysis (GSEA) revealed that transcripts encoding proteins belonging to the same pathway often show similar turnover dynamics. Furthermore, transcript isoforms show distinct dynamics suggesting that RNA turnover is important in regulating mRNA isoform choice. Finally, splicing across newly made transcripts appears to be cooperative with either all or none type splicing. These data sets generated as part of ENCODE4 illustrate the intricate and coordinated regulation of RNA dynamics in controlling gene expression to allow for the precise coordination of cellular functions.

## INTRODUCTION

Gene expression is regulated at many levels in mammalian cells. Transcription initiation is directed by epigenetic marks allowing regulatory DNA elements to be accessible to activating or repressive transcription factors. Epigenetic regulation to compact chromatin restrict transcription and we recently reported that only about 20% of the genome in human cells is actively producing measurable amounts of RNA^1^. Furthermore, chromatin organization dictates the proximity of genes to regulatory elements^2–5^. The abundance and quality of synthesized RNAs are then regulated through different processing and degradation pathways in the nucleus and the cytoplasm. For example, the nuclear RNA exosome participates in the processing and degradation of precursors of tRNA, rRNA and mRNA as well as RNA generated from enhancer elements (eRNA) and promoter upstream regions (PROMPTs)^6–10^. Furthermore, RNA degradation may take place in the cytoplasm in association with translation via nonsense-mediated decay (NMD)^11–14^, nonstop-mediated decay (NSD)^15, 16^ or no-go-decay (NGD)^17^.

RNA turnover can be assessed by measuring the decay of steady-state RNA either after transcriptional inhibition or following a chase of metabolically bromouridine-labeled steady-state RNA^18–20^. While transcription inhibition with pharmacological inhibitors such as actinomycin D is stressful to cells, metabolic labeling of RNA with bromouridine for extended periods of time has minimal effects on transcription, post-transcriptional processing, as well as cell viability^20, 21^. In approach-to-equilibrium experiments, actively synthesizing RNAs in cells are labeled for increasing periods of time until the labeled population is at steady-state and decay rates can be determined by comparing labeled RNA abundance at different labeling times with the steady-state RNA levels^22^. Another popular approach is to label nascent RNA with a short pulse of a uridine analog and then compare the RNA abundance of the nascent transcriptome with the steady-state transcriptome. Such techniques include SLAM-seq^23^, Time-lapse-seq^24^ and TUC-seq^25^. Nano-ID can be used to assess isoform-specific turnover^26^ and Dyrec-seq employs double metabolic labeling with Bromouridine and 4sU for synthesis and degradation determinations^27^. Finally, INSPEcT is a computational approach using steady-state RNA-seq data to assess RNA turnover rates without the need of metabolic labeling^28^.

When comparing nascent RNA synthesis to steady-state RNA levels in cells it is assumed that RNA degradation follows first order exponential decay kinetics^29^. However, the kinetics of decay may be differentially affected by quality control processes removing aberrant RNA and by cytoplasmic degradation of mature mRNAs. To assess the dynamics of RNA without relying on steady-state RNA comparisons, we used BruChase-seq, which assesses the abundance of different RNAs at different time points after pulse-labeling with bromouridine and a uridine chase^30, 31^. By collecting RNA at different time points after labeling, transcriptomes of defined ages can be studied and compared to each other in an unbiased way^32^. In these studies, Bru-labeled RNA was collected either directly after labeling (nascent RNA) or after a 2 or 6-hour chase in uridine.

We recently reported, using Bru-seq, that co-transcriptional splicing is removing only about half of the introns from pre-mRNA in human cells^21^. Similar findings have been reported in studies subjecting nascent RNA to long-read RNA sequencing techniques where transcripts were found to be either fully spliced or not spliced at all, with a smaller group of transcripts showing a mixture of spliced and retained introns^33,34^. However, after a 6-hour chase time, the transcripts are almost completely spliced. This scenario suggests that the intron-containing transcripts are either removed from the transcriptome by quality control mechanisms and degradation^35^, or they are post-transcriptionally spliced^34^.

The results from our study using BruChase-seq across 16 human cell lines, show that RNA turnover plays a major role in regulating gene expression. The trajectories of degradation of RNA synthesized from individual genes were found to differ widely both during the early time interval (0-2h) and during the later time interval (2-6h). Furthermore, the pattern of transcript degradation appeared to be coordinated among genes coding for proteins belonging to the same cellular pathways as previously suggested^29, 32, 36–41^. Interestingly, genes coding for constituents of large molecular machines, such as the proteasome, spliceosome, ribosome, or the oxidative phosphorylation machinery, showed high rates of synthesis and turnover during the first 2-hour chase period. However, the transcripts that survived this initial purge were found to be very stable during the next time interval of chase suggesting a unique logic for regulating RNA turnover for these multi-protein complexes. Furthermore, assessment of turnover rates of different mRNA isoforms synthesized from the same gene showed different RNA degradation rates, suggesting that the choice of which isoform to express may not only be regulated during splicing but also by targeted post-transcriptional turnover. Finally, our results suggest that nascent pre-mRNAs are either fully spliced or not spliced at all and that the unspliced pool of transcripts rapidly diminishes. This data set generated for ENCODE4 gives us a comprehensive insight into the regulation of RNA processing in coordinating gene expression to accommodate homeostatic cell function.

## RESULTS

### Using BruChase-seq to assess RNA degradation dynamics across 16 cell lines

It is well known that certain RNA species are very unstable, such as introns, enhancer RNA (eRNA) and promoter upstream transcripts (PROMPTs) produced upstream of most active gene promoters in the divergent direction of the gene^42^. The functions of eRNA and PROMPTs are not well characterized but these transcripts are short (<4 kb) and are turned over very rapidly by the RNA exosome^8, 43^ making it difficult to capture them from steady-state RNA isolations. RNA is also generated from transcriptional read-through at the ends of genes^1, 44^ as well as from repetitive DNA sequences, which make up over half of the human genome^45^. To assess the relative turnover of these different classes of RNA, we performed nascent RNA Bru-seq and BruChase-seq. Newly labeled RNA were captured at time zero (Bru-seq or 0h) or after a 2-hour or a 6-hour uridine chase interval (BruChase-seq, 2h and 6h) across 16 human cell lines **(Fig. 1a**). These experiments were performed on two independent growths of these cell lines and the RNA was sequenced to a depth of ∼50 million paired-end reads^1^ (**Table S1**). As RNA ages, it is expected that the fraction of intronic signal should decrease as splicing and degradation take place while the fraction of exonic signal should increase. Indeed, compared to the 0h samples, the exonic read fraction across all 16 cell lines increased 2.9-fold for the 2h sample and about 4.3-fold for the 6h sample while intronic reads decreased by 0.66-fold and 0.49-fold for 2h and 6h, respectively. We next compared the stability of introns relative to exons over the two time periods. The results show that compared to the stability of exons, intronic stability was only 0.23-fold and 0.11-fold after a 2-hour and a 6-hour chase period, respectively (**Fig. 1b, Table S2**). PROMPT RNA and eRNA showed similar degradation patterns to intronic sequences while transcription readthrough RNA was slightly more stable. Interestingly, RNA generated from repetitive sequences showed an extremely fast turnover rate. It is possible that this type of RNA is specifically targeted for rapid degradation because its presence can elicit strong inflammatory signaling^46^. These results show that different types of RNA species are degraded by different kinetics.

**Figure 1.**
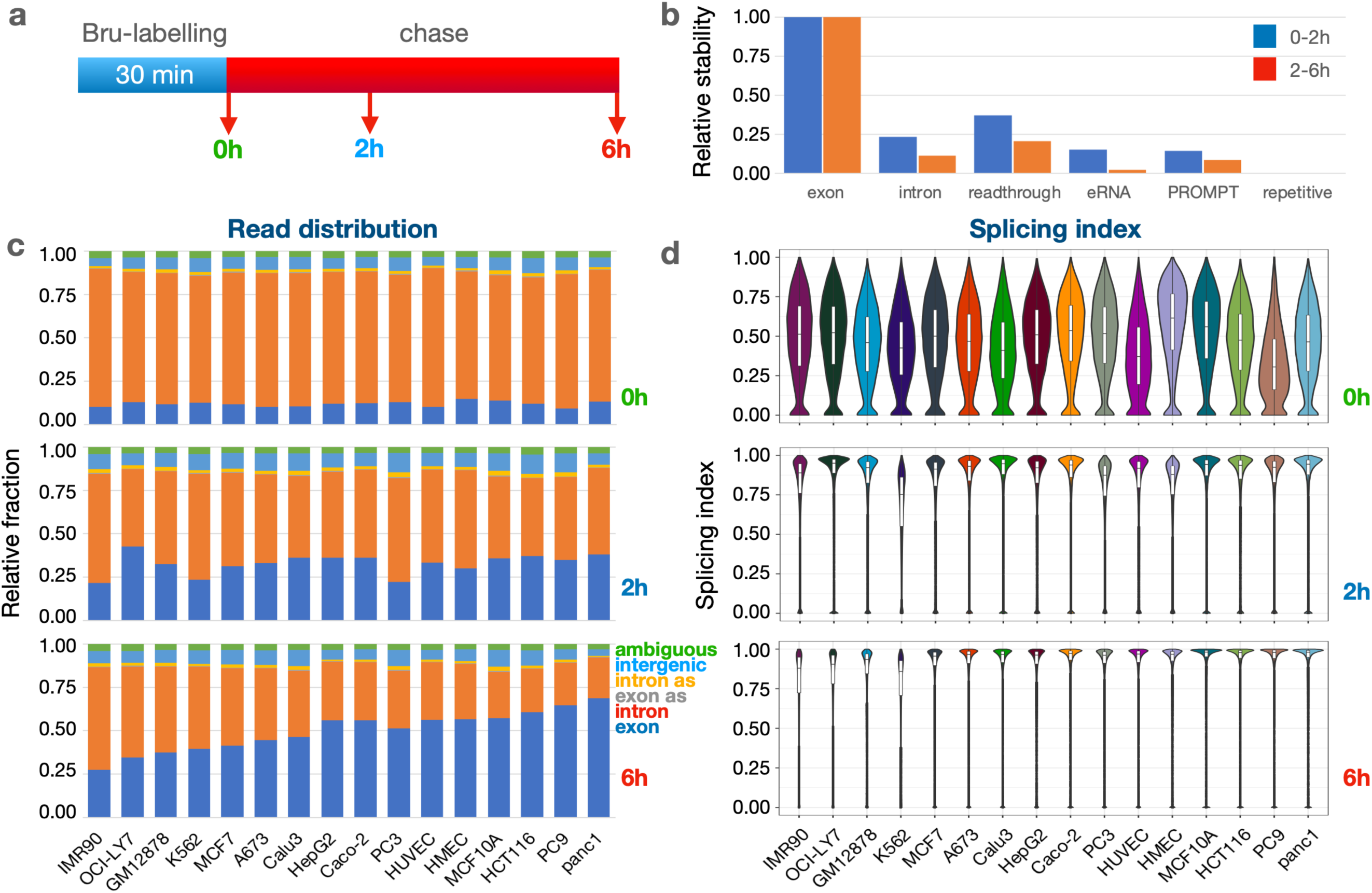
**a**) Experimental outline. **b**) Relative stability of different RNA categories compared to protein-coding exons. Data are averages of values collected from 16 human cell lines (Table S1). Relative stability at 0h (green), 2h (blue) and 6h (red). **c**) Read distribution of different RNA categories at 0h (top), 2h (middle) and 6h (bottom) across 16 human cell lines. Exons (dark blue), introns (red), exons antisense (orange), intergenic RNA (blue) and ambiguous (green). **d**) Splicing index (SI) across 16 human cell lines at 0h, 2h and 6h after Bru-labeling. The SI values across 24272 introns are displayed in violin plots showing the median (hatchbar) and the 95^th^ percentile of the data in a white boxplot.

We next compared the fraction of reads obtained from all exons, introns, exon antisense, intron antisense, intergenic sequences across the 16 cell lines. In addition, we included ambiguous reads which are reads that could not be mapped. At time 0h, the intronic signal made up 72-80% of the total signal and exonic reads made up 9-13% across the cell lines, while intergenic (or non-annotated) reads made up 5-9% and ambiguous reads made up 3-4% of all the reads. (**Fig. 1c, Table S3**). Antisense RNA amounted to less than 2% of all the reads. Following a 2h and 6h chase, exonic reads increased and intronic reads decreased while the fractions of antisense, intergenic and ambiguous reads stayed fairly constant. We noticed a large discrepancy in the intronic signal across the 16 cell lines at the later time point. For example, a much higher retention of intronic signal following a 6-hour chase was observed in IMR90 cells (61%) than in Panc1 cells (28%). This may suggest that either the rate of splicing or the rate of intron turnover differ between the cell lines.

To explore whether the high retention of intron signal in IMR90 may correspond to poor splicing, we estimated the splicing index (SI) of genes that were expressed across all 16 cell lines as previously described^21^. Co-transcriptional SI, defined as a measure of splicing occurring during the 30-minute bromouridine-labeling period (0h), showed that the median SI values ranged from 0.30 (for PC9) to 0.61 (for HMEC) (**Fig. 1D, Table S4**). Upon assaying the RNA two hours after labeling (2h), the median SI values jumped to 0.75 for K562 cells and to around 0.87 or higher for most of the other cell lines. Finally, median SI values increased to over 0.9 following a 6-hour chase for most cell lines. These results show that, on average, only about half of the splicing junctions flanking introns are co-transcriptionally spliced but with time, splicing of the transcriptome becomes more complete. This may be the result of either post-transcriptional splicing activities or the RNA degradation machinery purging the transcriptome of intron-containing transcripts. Indeed, it has been shown using the BruChase-seq technology that intron-containing transcripts are considerably less stable than spliced transcripts^35^, suggesting a targeted degradation of unspliced transcripts.

The high retention of introns in IMR90 cells after a 6-hour chase period (61%) does not correlate to a conversely low splicing index (>87%), suggesting that these introns are spliced out but not efficiently degraded during the chase period in this cell line. We do observe a slight trend that the cell lines with highest intron retention were also the cell lines with the lowest splicing indexes at 2h and 6h.

### Cell type and pathway-specific RNA dynamics

To compare the trajectory of Bru-labeled RNA degradation over time across the 16 cell lines, we obtained relative stabilities of expressed genes in the early (0-2h) and late (2-6h) chase periods. We used only exonic reads for this measurement as introns are rapidly turned over post-splicing with kinetics very different from the exons in the mature mRNA (see Fig. 1b). We found that correlations of gene stabilities among the cell lines were generally stronger at 2h than at 6h chase and that hierarchical clustering was quite different between the two chase periods (**Fig. 2a-b**).

**Figure 2.**
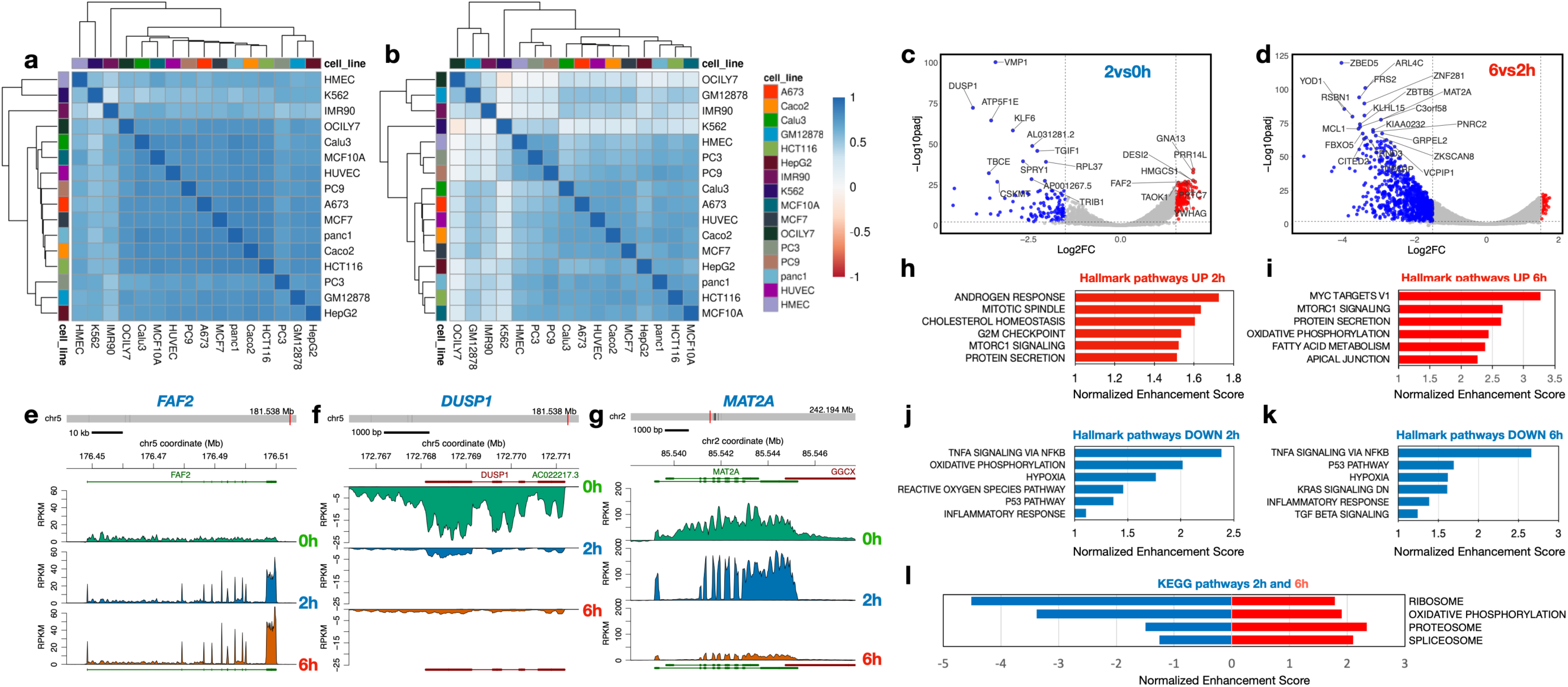
Gene and pathway-specific relative RNA stability. **a**) We used pairwise correlations (Pearson’s r) to cluster the cell lines according to their relative stability of transcripts from all expressed genes during the 0-2h time period **b**) and during the 2-6 h time period. **c**) Volcano plot of relative stability of transcripts 0-2h in Panc1 cells. **d**) Volcano plot of relative stability of transcripts 2-6h in Panc1 cells. **e**) Read coverage profiles for FAF2, **f**) DUSP1 and **g**) MATA2 in Panc1 cells at 0h, 2h and 6h. GSEA analyses of relative RNA stability in Panc1 cells showing up-regulated Hallmark pathways at 2h **h**), and at 6h **i**) and Hallmark pathways down-regulated at 2h **j**) and at 6h **k**). **i**) GSEA analyses in Panc1 cells of 4 GSEA KEGG pathways involving multiprotein complexes showing down-regulated RNA stability at 2h but up-regulated RNA stability at 6h.

To learn more about whether turnover rates for protein-coding gene transcripts change over time, we compared the relative stability of individual genes during the early and late chase period. As an example for these analyses, we chose the Panc1 cell line which showed low intron retention and high splicing index at 6 h compared to other cell lines (**Fig. 1c&d**). Using an expression threshold of >0.5 RPKM in the 0h sample, we found that the relative stability increased significantly for 161 genes and decreased significantly for 114 genes in Panc1 cells during the first 2 h after synthesis (-Log10padj >2, log2FC >+/-1.5) (**Fig 2c, Table S5a&b**). For the 2 h to 6 h period, we found that the relative stabilities of 66 transcripts increased and 885 decreased significantly (**Fig. 2d, Table S3**). In many of the 16 cell lines there are more transcripts with higher relative turnover between 2 h and 6 h than between 0 h and 2 h (**Extended Fig. 1**). From the read coverage profile of *FAF2* (**Fig. 2e**) it can be seen that reads are distributed across the gene at 0h but concentrate to the exonic regions at 2h and 6h. This gene exemplifies genes which transcripts show high relative stability across both time periods (**Fig. 2e**). In contrast, *DUSP1* transcripts show low relative stability at both time points (**Fig. 2f**). Finally, *MAT2A* transcripts show high relative stability during the first 2 hours but then appear to be extensively degraded during the next 4-hour chase period (**Fig. 2g**). These findings show that degradation rates differ between transcripts and do not necessarily follow first order kinetics^47^.

To better understand what cellular processes may be influenced by relative gene stability, we performed gene set enrichment analysis (GSEA) using rlog fold-change for ordering of the gene lists^48^ (2h vs. 0h or 6h vs. 2h) to rank genes. For Panc1 cells, Hallmark gene sets such as “androgen response” and “mitotic spindle” were enriched for high-stability genes after the early period (0-2h) (**Fig. 2h**) and “MYC targets V1” and “protein secretion” after the late period (2-6h) (**Fig. 2i**). Gene sets enriched in low-stability genes included “TNF signaling via NFkB” and “oxidative phosphorylation” for the early period (**Fig. 2j**) and “TNF signaling via NFkB” and “p53 pathway” for the late period (**Fig. 2k**). Interestingly, genes in KEGG pathways such as “ribosome”, “oxidative phosphorylation” (mitochondria), “proteasome” and “spliceosome” produced transcripts that were relatively unstable at the early period but were of above average stability at the late period. (**Fig. 2l**). This trend holds up when analyzing the stability of transcripts of these pathways across all 16 cell lines (**Extended Fig, 2a**). These genes code for large biomolecular machines and it is possible that the cellular strategy is to over-produce these transcripts and then rapidly “purge” the excess to arrive at stoichiometric levels for the biosynthesis of these machines.

A number of Hallmark gene sets were commonly enriched for transcript stability (both positively and negatively) across the 16 cell lines at the early period (0-2h) (**Extended Fig. 2b**). High gene stability examples include “androgren response” (NES > 1 in all cell lines) and “mitotic spindle” (NES > 1 in 15/16 cell lines) and low gene stability examples include “TNF signaling via NFkB” (NES < −1 in 14/16 cell lines) and “oxidative phosphorylation” (NES < −1 in 15/16 cell lines). Transcripts with high relative stability during the 2-6h chase included “MYC targets” and “oxidative phosphorylation” while “TNFA signaling” and “hypoxia” gene sets are associated with high turnover. For KEGG pathways, we observed a similar trend that the degradation of transcripts belonging to similar pathways were coordinated (**Extended Fig. 2c**). These results suggest that transcripts coding for proteins belonging to a specific pathway share a similar turnover rate and that this logic is consistent across cell lines.

### Post-transcriptional regulation of mitochondrial encoded transcripts

The circular mitochondrial genome is transcribed as three polycistronic transcripts covering 2 rRNA genes, 22 tRNA genes and 13 protein-coding genes needed for the mitochondrial oxidative phosphorylation system (OXPHOS)^49^ These polycistronic transcripts are processed into individual transcripts via the “tRNA punctuation model”^50^, although additional cleavage sites in these transcripts have been observed ^51^. It has been noted that the levels of individual mRNAs are regulated by differential degradation^52^ and that Suv3p helicase plays a role in regulating processing and turnover of mitochondrial transcripts^53^. When exploring the Bru-seq and BruChase-seq data from the 16 cell lines we observe that the expression of the mitochondrial genes is differentially regulated post-transcriptionally. It can be seen from the distribution of read counts across the mitochondrial genome that *ND1*, *ND2* and *ND3* genes produce relatively unstable transcripts while *CO1*, *CO3*, *ATP6*, *ATP8* and *ND4* genes generate relatively stable transcripts (**Extended Fig. 3**). However, K562 and PC3 cells show higher stability of the *ND2* transcript compared to all other cell lines. In concurrence with a previous study, we notice the expression of antisense RNA for the *ND5* gene^51^, which is a very unstable transcript except in the A673, Caco2, HUVEC and K562 cells where this ncRNA was relatively stable (**Extended Fig. 3**). Thus, our data suggest both gene- and cell type-specific post-transcriptional regulation of mitochondrial gene expression.

**Figure 3.**
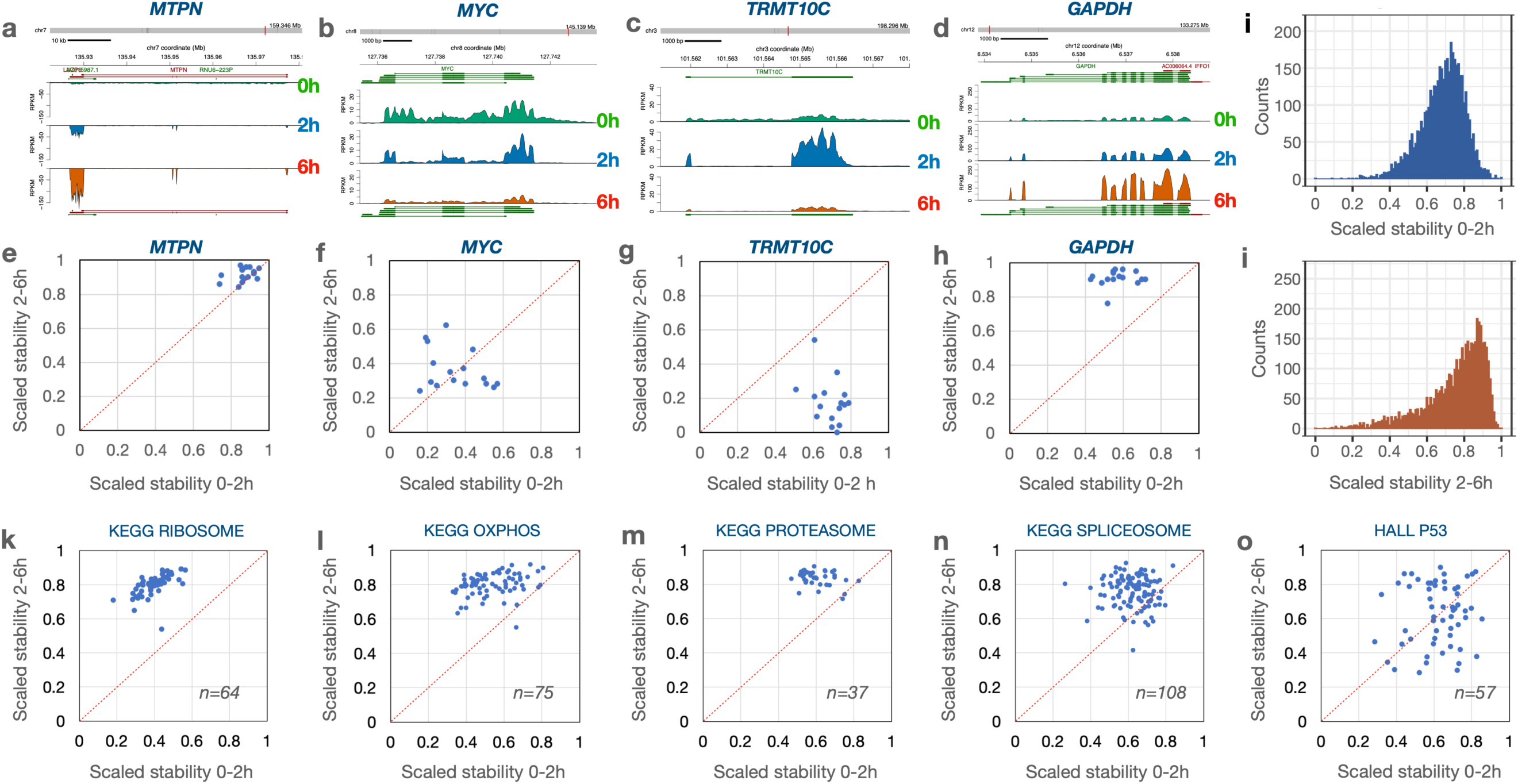
Different RNA stability dynamics for selected genes. **a-d**) Graphs displaying read counts across selected genes in Panc1 cells at the three time points. **e-h**) The log2FC scaled stability scores of the transcripts portrayed in a-d were plotted for each of the 16 cell lines in a 2-dimensional matrix. **i**) Distribution of scaled stability scores of all transcripts in Panc1 cells during the 0-2h time period and **j**) during the 2-6h period. **k-n**) Distribution of scaled stability scores for transcripts belonging to selected GSEA pathways. The stability scores of each individual transcript were averaged across the 16 cell lines and plotted in a 2-dimensional matrix with the number of genes sampled in the pathway indicated.

### Unique RNA dynamics for individual genes

We next compared the dynamics of degradation for transcripts of individual genes across the 16 cell lines. In **Figure 3a-d** we plotted read coverages for four different genes in Panc1 cells with widely different dynamics to display differences in relative degradation. To compare the trajectories of degradation of transcripts of individual genes during the two chase periods across the 16 cell lines, we scaled the log2FC relative stability values between 0 and 1 (**Table S6**). The result of this scaling for the *MTPN* transcripts shows that data points from all 16 cell lines clustered in the upper right corner implicating high stability across both time intervals (**Fig. 3e**). In contrast, transcripts from the *MYC* gene shows a high degree of degradation during both time periods across all 16 cell lines (**Fig. 3f**). Both *MTPN* and *MYC* have transcripts that center around the diagonal, implying that the degradation rates were comparable during both time intervals. In contrast, most of the data points for the *TRMT10C* transcript fell below the diagonal (**Fig. 3g**) reflecting initial high stability followed by low stability while the scaled stability values for *GAPDH* fell above the diagonal line, implicating initial fast turnover followed by high stability (**Fig. 3h**). These results show that some transcripts have different degradation rates during the first 2 hours and the next 4 hours following synthesis and thus, suggest that not all transcripts follow first order degradation kinetics^47^.

The patterns of transcript stability were in general similar across the cell lines, implicating that the transcript, rather than the cell type, is the major determinator of these degradation patterns. However, we found exceptions where transcripts showed cell type specific RNA dynamics trajectories (**Extended Fig. 4**). The distribution of the scaled RNA stability values of all expressed genes in Panc1 cells are shown for both the first 2 h (**Fig. 3i**) and the next 4 h (**Fig. 3j**). Similar analyses for all 16 cell lines can be found in **Extended Fig. 5**.

**Figure 4.**
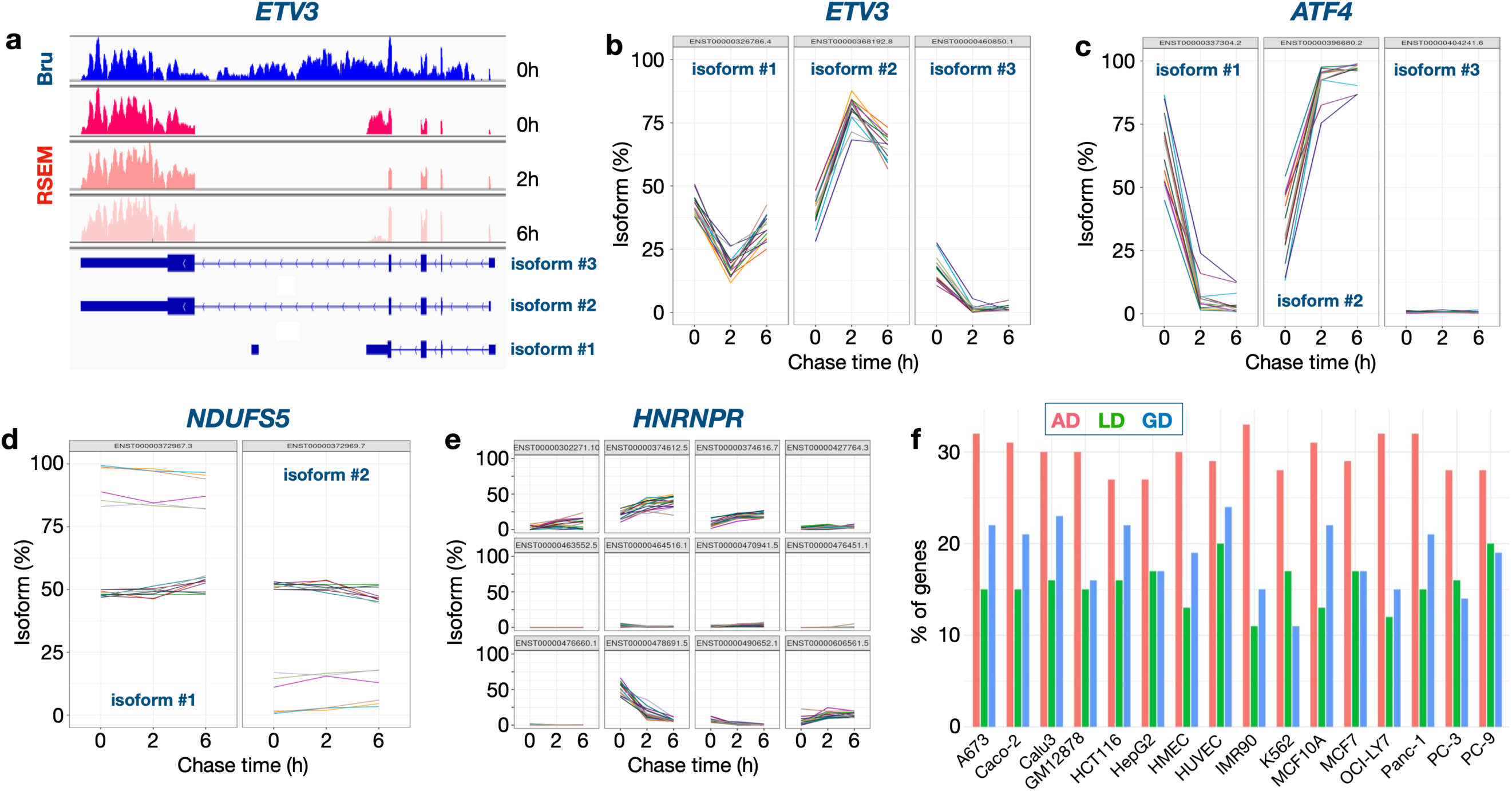
Isoform-specific post-transcriptional processing. a) Coverage of captured Bru-labeled RNA across ETV3 isoforms. Isoform percentages as computed by the RSEM pipeline for transcript isoforms generated by ETV3 (b), ATF4 (c), NDUFS5 (d) and HNRNPR (e) genes for the tree time points. (f) Percentage of genes (1076 genes with at least 2 isoforms) under study whose isoforms showed varying patterns of dominance over time: always dominant (AD), gain dominance (GD) and lose dominance (LD).

**Figure 5.**
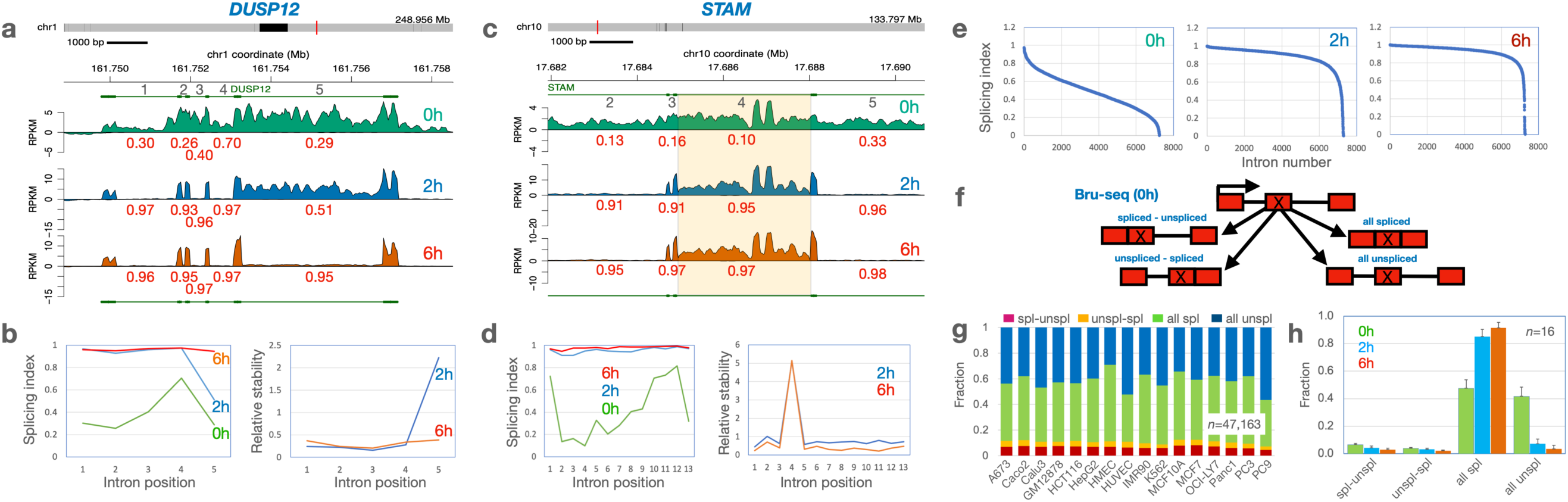
Splicing patterns differ within and between transcripts. **a**) Trace diagrams of RPKM values across the DUSP1 gene in A673 cells at 0 h, 2 h, and 6 h with red numbers indicating SI values for individual introns numbered in black. While introns 1-4 show >93% splicing at 2 h, intron 5 only shows ∼50% splicing but approaches complete splicing by 6 h. **b**) SI values for the introns along the DUSP12 transcript (left graph) and the corresponding relative intron abundances (right) are shown for the different time points. **c**) Trace diagram of RPKM values across the introns 2-5 of the STAM gene in A673 cells showing retention of intron 4 throughout the time course despite being efficiently spliced at 2 h. **d**) SI values along the STAM transcript (left) and relative intron abundance (right) in A673 cells. **e**) distribution of SI values for 7231 introns from 626 protein-coding genes at 0 h, 3 h and 6 h. (**Table S4**). **f**) Diagram of the formation of four different splicing patterns. **g**) Fraction counts of the four co-transcriptional splicing pattern bins for 47,163 introns across 16 cell lines. **h**) Fraction counts of averages from each of the 16 cell lines for the different time points.

We next displayed the coordinates for normalized stabilities of transcript coding for proteins belonging to the same functional pathways. For each transcript we averaged the 2h vs 0h and the 6h vs 2h stability values from all 16 cell lines. For the 64 transcripts expressed across all 16 cell lines representing the GSEA “KEGG ribosome” pathway, the trajectories of degradation are different between the two time points in that the initial phase show a higher rate of degradation than during the later period (**Fig. 3k**). Similar patterns were observed also for transcripts representing the pathways “oxidative phosphorylation”, “proteasome” and “spliceosome” (**Fig. 3l-m**). These findings are in concordance with the data presented in **Figure 2l** and in **Extended Fig. 2c**.

Furthermore, breaking out specific genes from these pathways, we observe high concordance across the 16 cell lines in that individual transcripts show higher degradation during the first 2-hour chase than during the next 4 hours (**Extended Fig. 6**). For most other pathways analyzed in this way, we find distributions that are centered around the diagonal suggesting a constant degradation rate over both time periods. For some pathways, such as the p53 pathway (Fig. 3o), the stability scores of individual transcripts show a wider distribution suggesting that there was less pathway-specific regulation of post-transcriptional processing of transcripts in this pathway (**Fig. S7**).

These data suggest that transcript stability most often is transcript specific with similar dynamics across the cell lines although some cell-type specific regulation was found for a smaller set of transcripts.

### Isoform-specific post-transcriptional processing

Most human genes generate multiple transcript isoforms due to the use of alternative promoters, splice junctions or poly(A) sites^54^. While the choice of a specific promoter for transcriptional initiation is regulated by transcription factor binding and epigenetic marks^55–57^ the choice of splicing junctions is regulated by protein binding to splicing enhancer and silencer sequences in the RNA molecule^54^. What is less understood is whether isoform-selective RNA turnover may contribute to the emergence of a dominant isoform from a field of synthesized isoforms. To determine whether isoform-selective post-transcriptional processing operates in human cells, we analyzed the Bru-seq (0h) and BruChase-seq (2h & 6h) data using the RSEM software tool^58^. We expressed the isoform abundances as a percentage of all isoforms for each gene and time point. Both *ETV3* and *ATF4* genes (**Figs 4a-c**) show opposing abundance patterns for their respective isoforms across a majority of the 16 cell lines, with one isoform’s abundance decreasing and the other’s increasing over the time course. *NDUFS5* (**Fig. 4d**) isoforms abundances vary between cell lines. While both isoforms of the gene seem to coexist equally throughout the time course in 10 cell lines, isoform #1 is the dominant isoform in the remaining 6 cell lines. Its dominance is established within the first 30 mins of production (0h) and is maintained through the chase periods. *HNRNPR* (**Fig. 4e**) shows a behavior similar to *ETV3* and *ATF4* genes with its 3 isoforms (#2, 3 and 5). Taking an isoform percentage of 50% as the threshold for annotating an isoform as dominant (or major), roughly 30% of the genes we studied showed a pattern wherein one of their isoforms established dominance at 0h and stayed dominant through the time course (always dominant, AD). Examples include isoform #1 in NDUFS5 in 6 cell lines. Around 20% of the genes had an isoform that gained dominance (GD, *ETV3* and *ATF4* isoform #2), starting below 50% prevalence but moving above during the time course, while roughly 15% of the genes had an isoform that lost dominance (LD) over time. *ETV3* and *ATF4* would be genes that would get classified under both GD and LD. The abundance patterns for the remaining genes could not be clearly delineated. These results suggest that many genes generate multiple isoforms up front that are then regulated by isoform-specific turnover to arrive at the desired stable isoform.

### Splicing dynamics and “all or none” splicing

Introns are spliced co-transcriptionally from nascent RNA as it exits the RNA polymerase^59, 60^. Thus, it would be expected that introns are removed in a sequential manner as the RNA molecules exit the RNA polymerase. However, several studies have reported that the order of splicing does not always proceed in the direction of transcription ^34^ and that co-transcriptional splicing is often not complete^21^. To explore whether transcripts show consistent SI values for individual introns across the transcripts or whether each intron of a transcript is removed independently of each other, we analyzed multi-intron genes producing a single annotated isoform (RPKM >=0.5 at 0h, n=626 genes with 7231 introns).

When analyzing the SI values of individual multi-intron transcripts, we found that the splicing efficiency along the transcripts differed markedly. Transcripts synthesized from the *DUSP12* gene in the A673 cell line shows that most introns are still present following the 30-min Bru-labeling of the nascent RNA and they have not been spliced out evenly (**Fig. 5a**). Following a 2-hour chase period, introns 1-4 are almost completely spliced out (>93%) and the introns are turned over. In contrast, intron #5 is poorly spliced even at the 2-hour time point but is efficiently spliced out and degraded by 6h (**Fig. 5b**). This example suggests that co-transcriptional splicing is not coordinated across all introns of this transcript and that one intron is initially retained in the transcript due to incomplete splicing. Intron # 4 in the STAM gene, on the other hand, is spliced out by the 2h chase time (**Fig 5c-d**) yet remains present even in the later chase timepoint. A similar pattern is observed for this intron across all 16 cell lines (**Extended Fig. 8 a&b**). It is possible that after splicing, this intron serves a non-coding function in the cells and is therefore not targeted for immediate degradation. Finally, we compared the distribution of splicing values of all the 7231 introns at the three different time points and as expected, the distribution of SI values increased to represent more spliced introns over time (**Fig. 5e**). These results illustrate the complexity of the splicing process and that the splicing index increases over time as a result of either post-transcriptional splicing or purging of unspliced transcripts.

It has been shown using long read sequencing that a large portion of transcripts show an “all or none” splicing pattern^33, 34, 61^. Since the sequencing of the Bru-seq assays were performed using 150 nucleotide paired-end sequencing of fragments 400-500 bp in length, it was possible to use this data to assess two splicing events on the same sequenced fragment. While still focusing on single-isoform genes (n=4578 genes with 47,163 exons), we computed the splicing patterns of fragments spanning an exon (instead of an intron as studied earlier). The various splicing possibilities (**Fig. 5f**), as captured by the sequencing reads were computed across each exon and summed up.

The two biggest groups of splicing patterns that emerged were “all spliced” and “all unspliced” in all 16 cell lines (**Fig. 5g, Table S7**). With chase, the “all spliced” fraction increased at the expense of the “all unspliced” fraction (**Fig. 5h**). We also observe that of the two smaller groups depicting intermediate splicing patterns (spliced-unspliced and unspliced-spliced), about twice as many exons with the upstream intron only spliced were found compared to exons with the downstream intron only spliced. Taken together, these findings show that there are two major modes of transcription; one that leads to complete splicing and one that leads to no splicing. Our study concurs with, and extends, earlier findings^33, 34, 61^ by showing that this pattern holds true across 16 human cell lines. Furthermore, we show that the fraction of unspliced transcripts rapidly diminishes over time. Whether this increase in splicing index is due to post-transcriptional splicing or targeted degradation of the pool of unspliced transcripts will need to be further elucidated in future studies.

## DISCUSSION

RNA turnover is a critical component in the regulation of gene expression. While substantial attention has been given to the mechanisms of transcription initiation via transcription factors and epigenetic modifications, less is known about how RNA turnover is regulated^62^. In these studies, we used Bru-seq and BruChase-seq technologies to assess nascent RNA synthesis, splicing and degradation. BruChase-seq differs from other technologies that have been used to assess RNA degradation in that it does not depend on comparisons with steady-state RNA levels. Instead, Bru-labeled RNA is monitored as it is aging during uridine chase periods (2h or 6h in this study) and then captured and relative RNA abundance is assessed genome-wide. While other studies assessments of steady-state cellular RNA have shown that degradation is occurring by first order kinetics, our BruChase-seq approach found clear exceptions to this. Performed across 16 human cell lines as part of the ENCODE 4 project we observed that several transcripts showed different degradation kinetics during the two time periods. Furthermore, we found that RNA degradation varies widely among different transcripts, is mostly transcript dependent with only a few transcripts showing cell type-specific regulation.

As expected, intronic sequences showed a rapid turnover compared to exonic sequences (**Fig. 1b**). The same was true for readthrough transcripts, PROMPT RNA and eRNA. RNA generated from repetitive sequences were highly unstable which may be by design since this RNA can elicit an inflammatory response if not rapidly eliminated^63^. The rapid turnover of the vast amount of intronic RNA is critical for recycling of ribonucleotides that then can take part in new synthesis. Interestingly, we found that cell lines differed quite a bit in the turnover of intronic sequences where the normal fibroblasts IMR90 retained over 60% of the intronic sequences after 6h of chase (**Fig. 1c**). This high intron retention in IMR90 was not due to lack of splicing but rather must be the result of slow degradation of spliced introns. Co-transcriptional splicing rates were found to be between 30% and 60% across the 16 cell lines but the splicing index increased to near 90% after a 2h chase (**Fig. 1d**).

It has been previously suggested that there is a physiological logic governing the decay rates of functionally linked transcripts^29, 32, 36–41^. This is found for transcripts encoding multimeric protein complexes both in yeast^41^ and in human cells^36^. Furthermore, transcripts linked to regulatory functions, such as transcription factors, typically have short half-lives so that cells can rapidly respond to environmental stimuli and challenges^32, 37, 39^. This was found for some but not all transcripts coding for transcription factors where *KLF6*, *KLF10* and *MYC* were very unstable transcripts while *TP53*, *HIF1A* and *SMAD5* are examples of rather stable transcripts (Flagship). We observed an unusual pattern of RNA degradation for transcripts encoding protein complexes such as the ribosome, the proteasome, the spliceosome and the oxidative phosphorylation machinery where the degradation patterns of these transcripts differed between the first 2-hour uridine chase and following 4-hours of chase where the initial degradation was extensive but reduced during the latter chase period. (**Figs. 3k-n, Extended Fig. 2c**). It is possible that the cellular strategy to obtain stoichiometric amounts of subunits for multimeric protein complexes is to synthesize these transcripts in excess and then rapidly purge this transcript pool to obtain a stoichiometric set of transcripts that are then used for translation.

Most genes in eukaryotic cells produce multiple transcript isoforms through alternative splicing or selection of alternative promoters. Alternative splicing can lead to frameshifting followed by rapid degradation or result in the synthesis of unique proteins leading to the diversification of the proteome^54, 60^. The mechanisms by which alternative splicing is regulated is not fully understood but specific sequences in exons and introns are thought to attract or block the binding of RNA-binding proteins involved in the splicing process^64^. Furthermore, epitranscriptomic modifications of pre-mRNA and sequence-mediated structural features can affect splice-site selection leading to generation of preferred transcript isoforms^60^. In addition, cells may generate a plethora of transcript isoforms from a gene and then remove unwanted isoforms through selective degradation. Using the BruChase-seq approach we observed several genes producing multiple isoforms up front followed by the ascent of one or many isoforms at the expense of other isoforms (**Fig. 4**). This finding suggests that a prominent mechanism of isoform selection is through selective post-transcriptional degradation of unwanted isoforms. Future studies are needed to further elucidate the mechanisms dictating this selective degradation.

Splicing is thought to occur directly after the RNA exits the RNA polymerase when the splice sites are exposed to the splicing machinery^59^. However, sequencing of nascent RNA has provided evidence that a high level of introns remains in transcripts long after the RNA polymerases have finished transcribing^21, 33, 34, 61^. It has also been noted that the order of splicing does not always correspond to the order of the introns in the transcript. Our previous study using Bru-seq technology corroborated these studies in that we found that only about half of all introns are co-transcriptionally spliced and the order of splicing is not always occurring in a linear fashion^21^. Here we extended these studies of splicing across 16 cell lines measuring splicing index across time. We found examples of introns that showed poor splicing even 2 hours after synthesis while neighboring introns were efficiently spliced (**Fig. 5b**). We also found introns that were efficiently spliced at 2 hours but the intronic signal remained (**Fig 5d**) suggesting that it was selectively retained perhaps because it provides a specific function for the cells.

We used pair-end sequencing of fragments size-selected around 500-600 bp in our Bru-seq and BruChase-seq studies. Since many exons are about 150 nucleotides long, we were able to explore two splicing events on the same molecules. Adding up splicing events for the four possible different combinations of splicing around these exons, we found that most molecules were either all-spliced or all-unspliced across all 16 cell lines (**Fig. 5g**). This finding agrees with previous studies using long-read sequencing of nascent RNA^33, 34, 61^. When assessing splicing over time, the fraction of all-spliced molecules increased at the expense of the all-unspliced fraction which diminished over time. One interpretation of this increase in splicing index over time is that these transcripts are spliced post-transcriptionally. Alternatively, all splicing may occur co-transcriptionally while transcripts that did not splice undergo a time-dependent degradation to eliminate them from the transcriptome. Why would cells expend time and energy transcribing genes into pre-mRNA that would not become spliced and doomed to undergo degradation? One possible explanation is that this excessive transcription is utilized for DNA damage scanning in a mutation-suppressing mechanism as previously proposed^65^.

These studies using Bru-seq and BruChase seq across 16 cell lines as part of ENCODE4, provides a deep data set of the synthesis and degradation of RNA in human cells. Furthermore, many complementary transcription and chromatin assays were performed on the same cell growths making it possible to directly compare data from many different assays. We show here examples of the complexity of RNA dynamics in human cells and the clear patterns of cellular logic in post-transcriptional regulation of gene expression. This is an understudied area, and future studies will certainly uncover the importance of the precise regulation of RNA dynamics for cellular homeostasis and how defects in this regulation can lead to human disease.

## MATERIALS AND METHODS

### Cell lines

The following human cell lines were used in this study: A673 (Ewing’s sarcoma), Caco2 (colorectal cancer), Calu-3 (non-small lung cancer), GM12878 (normal lymphoblastoid), HCT116 (colorectal cancer), HepG2 (liver cancer), HMEC (normal human mammary epithelial cells), HUVEC (normal human umbilical vein endothelial cells), IMR-90 (normal lung fibroblasts), K562 (leukemia), MCF-7 (breast cancer), MCF-10A (normal breast epithelial), OCI-LY7 (B-cell non-Hodgkin lymphoma), Panc-1 (pancreatic cancer), PC3 (prostate cancer) and PC9 (lung adenocarcinoma). These 16 cell lines were assayed as part of the ENCODE 4 Deeply Profiled Cell Lines collection and cultured as 2 independent growths as described^1^.

### Bromouridine labeling, chase, and library preparations

Cells were incubated in 2 mM bromouridine (MilliporeSigma) for 30 min to label nascent RNA and then immediately lysed (0h) or chased in 20 mM uridine (MilliporeSigma) for either 2 hours (2h) or 6 hours (6h) as previously described^30, 31^. Cells were then lysed in TRIzol reagent (Invitogen), total RNA isolated and a Bru-labeled RNA spike-in cocktail was added^66^. Bru-labeled RNA was then captured from 100 µg of total RNA using anti-BrdU antibodies (BD Biosciences). Stranded library preparation was carried out according to documents linked to each experiment on the ENCODE portal (encodeproject.com) and is also described in detail elsewhere^1^.

### Alignment parameters

Alignment was carried out as described elsewhere^1^. Briefly: BruChase-seq libraries are sequenced on the Illumina Novaseq platform (150bp paired-end reads) to a depth of approximately 50 million read pairs per sample. Raw reads were first trimmed using BBTools to remove adapter sequences and were then pre-aligned to the ribosomal RNA (rRNA) repeating unit (GenBank U13369.1), and the mitochondrial (chrM) and EBV genomes (from the hg38 analysis set) using Bowtie2^67^, where any reads aligning to these sequences were recorded then removed. Finally, STAR^68^ was used to perform the alignment of the remaining reads to the reference genome (GRCh38).

### RNA abundance calculation for features of interest

Uniquely mapping reads (MAPQ>=255) to intergenic regions, introns and exons on the sense or antisense strand, were counted using custom scripts, and their relative fraction to all uniquely mapping reads in the standard chromosomes were calculated. The fraction of reads that mapped to more than one gene in the standard chromosomes were assigned to the ambiguous class. PROMPTs, intergenic enhancer regions, and RT segments were obtained similarly to those described elsewhere^1^. Briefly, intergenic enhancers were obtained from BruUV-seq peak calls that were intersected with the gencode v29 gene annotation (bedtools intersect -v) to remove those overlapping annotated genes. Intergenic RT segments were extracted using classifications defined by McShane et. al.^1^ where segments were retained if they were tagged as classes Ia or Ib, meaning they did not overlap a gene on the same strand. Counts for genes, PROMPTs, eRNA and read-through regions were obtained by calculating a fractional coverage of sequenced reads over each base in a strand-specific manner, and then summing the fractional coverages along the entire feature. Counts for repetitive elements were calculated using the fractional counting method implemented in the RepEnrich computational pipeline^69^.

### Splicing Index calculation

Splicing Index calculations were performed as previously described^21^. Briefly, an intron-centric Splicing Index (SI) was calculated for all introns common across the 16 cell lines in both Bru-seq and BruChase-seq assays, after merging the reads from the two replicates per cell line per assay. The SI was defined as a/(a+(b+c)/2), where a = split reads (i.e reads from exon to exon), b = 5 ‘Exon-Intron junction reads (with a minimum 10 bp overlap on either side of the junction), c = 3 ‘Intron-Exon junction reads (with a minimum 10 bp overlap on either side of the junction) and jrsum = junction reads sum (a+b+c).

### Dynamics of protein-coding RNA

Reads were counted in exons only ("exonic") per gene across Bru-seq (0h) and BruChase-seq (2h and 6h) libraries. For each cell line, counts from 2 replicates for each condition were normalized and compared using DESeq2 (DESeq2 v1.18.1, R v3.4.3)^48^.

The resulting log2 fold change values were used to represent stability between the 0 to 2 h and 2 to 6 h time points. Those genes not rejected by independent filtering in any time point comparison (DESeq2) for all 16 cell lines were retained (17049 genes for exonic stabilities). Pairwise correlations (Pearson’s *r*) were calculated and used to cluster the cell lines.

To compare the trajectories of degradation of transcripts from individual genes during the two chase periods across the 16 cell lines, we computed DESeq2-based log2 fold change (log2FC) values from two biological replicates using exonic reads using DESeq2^48^. Two separate log2FC values were computed, one based on 0h vs 2h timepoints, and the other on 2h vs 6h timepoints. We applied an expression filter of 0.5 RPKM in the earlier time point for both comparisons (at 0h for the 0h vs 2h comparison, and at 2h for the 2h vs 6h comparison) to ensure genes with a healthy expression were used as the starting point. Volcano plots were generated using log2FC and −Log10padj significance data from 2 biological experiments using VolcoNoseR^70^. Gene set enrichment analysis (GSEA 4.2.3.) were performed using a rlogFC ranked lists to obtain Hallmark and KEGG pathway gene set enrichments.

To compare the trajectories of RNA stability for genes across cell lines, we selected genes expressed at RPKM>0.5 for the earlier time point in the comparison across all 16 cell lines and we obtained 4415 genes that qualified after employing these strategies. We then scaled (using rescale function in R) the log2FC values between 0 and 1 per cell line for these 4415 genes, with 1 representing highest stability and 0 lowest stability. This allowed us to compare the RNA dynamics for 4415 genes across the 16 cell lines. The scaled coordinates for RNA stability for the two time periods were then plotted in a two-dimensional matrix.

### Isoform-specific RNA dynamics

For gene isoforms-specific analysis, we merged the raw data for both replicates of a cell line for each of the 3 assays (Bru-seq 0h, BruChase-seq 2h and BruChase-seq 6h) and processed them through the RSEM pipeline^58^. Specifically, merged reads were first trimmed using bbduk (bbTools v38.46 https://sourceforge.net/projects/bbmap/). Reads not rejected during trimming were aligned to the premap reference set (human rRNA, human mitochondrial genome, EBV genome and select spike-in references (*D. melanogaster* (dm6) rRNA, dm6 mitochondrial genome)). Reads that did not align to premap references were aligned to the main (genome) reference set (hg 38 GENCODE 29, select spike-in references (dm6 genome, *E. coli* genome (K-12 MG1655) and in vitro transcribed *A. thaliana* RNA sequences)) using the following RSEM options (rsem-calculate-expression --num-threads 16 --star --star-gzipped-read-file --output-genome-bam --sort-bam-by-coordinate --sort-bam-memory-per-thread 8G --temporary-folder/temp --keep-intermediate-files --time --paired-end). The RSEM output yielded isoform TPM and isoform percentage quantification files. For downstream analysis, we shortlisted genes that were expressed at >= 1 TPM in only the Bru-seq (0h) timepoint and common across all 16 cell lines which resulted in 1076 genes with at least 2 isoforms. An isoform with isoform percentage >=50% at a given timepoint was called the dominant or major isoform. A <u>shiny tool</u> was created that projects the isoform percentages for these common genes over the time course.

## Supporting information

Table S1 Sample Stats

Table S2-Relative Stability

Table S3-Read Distribution

Table S4-SI values 0h, 2h, 6h-final

Table S5a-RelativeStabilityPanc1-2h

Table S5b-RelativeStabilityPanc1-6h

Table S6-ScaledStability 2h & 6h

Table S7-all or none splicing

## ACKNOWLEDGEMENTS

We thank the personnel at University of Michigan Advanced Genomics Core for professional technical assistance and Manhong Dai and Fan Meng for administration and maintenance of the University of Michigan Molecular and Behavioral Neuroscience Institute (MBNI) computing cluster. This work was supported by grants from NHGRI (UM1 HG009382) and the Rogel Cancer Center Support Grant (P30 CA046592) by the use of the following Cancer Center Shared Resource(s): Cancer Data Science.

---

**Extended Figure 1.**
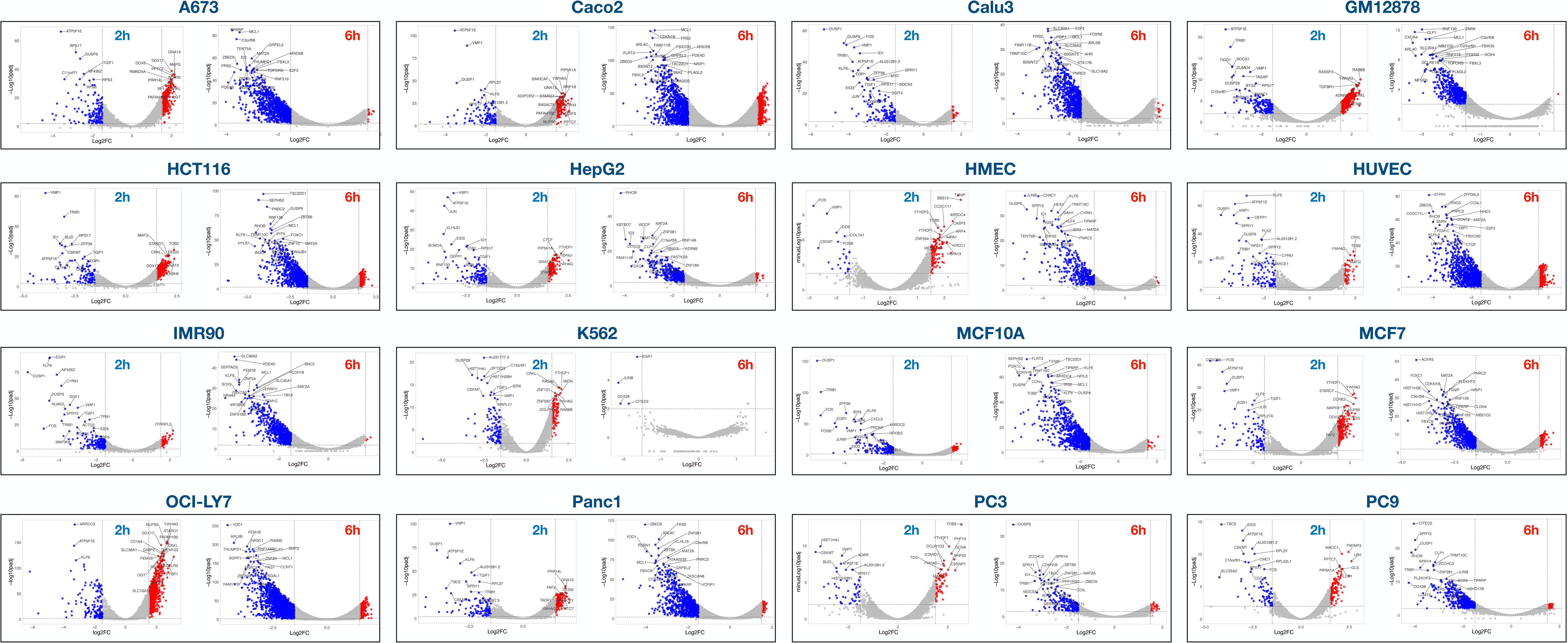
Volcano plots of relative RNA stability displayed as Log2FC vs −Log10padj for 0-2 h (left) and 2-6 h (right) time periods across 16 cell lines. Red dots represents transcripts with significantly high relative stability while blue dots represents transcripts with significant low relative stability.

**Extended Figure 2a.**
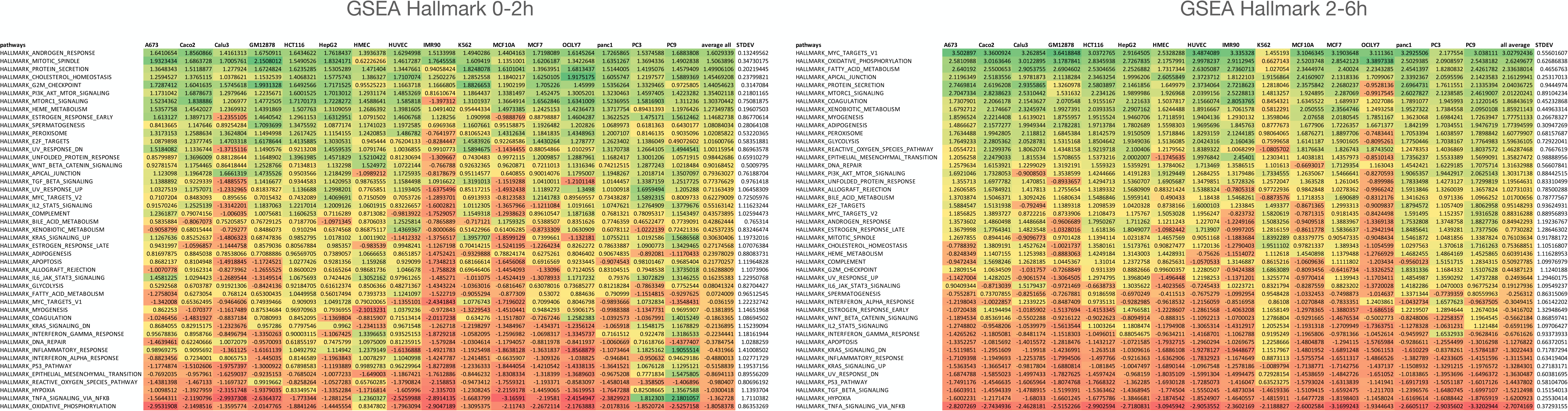
Co-regulation of RNA stability for transcripts with similar functions. GSEA analysis showing the Normalized Enhancement Score (NES) of relative stability of transcripts across 16 cell lines at 0-2h (left) and 2-6h (right). The different Hallmark pathways (rows) were averaged for all 16 cell lines and this average was used to order the different pathways from the most positive (green) to the most negative (red). The standard deviation of averages of the NES values of the 16 cell lines are shown in the column on the right.

**Extended Figure 2b.**
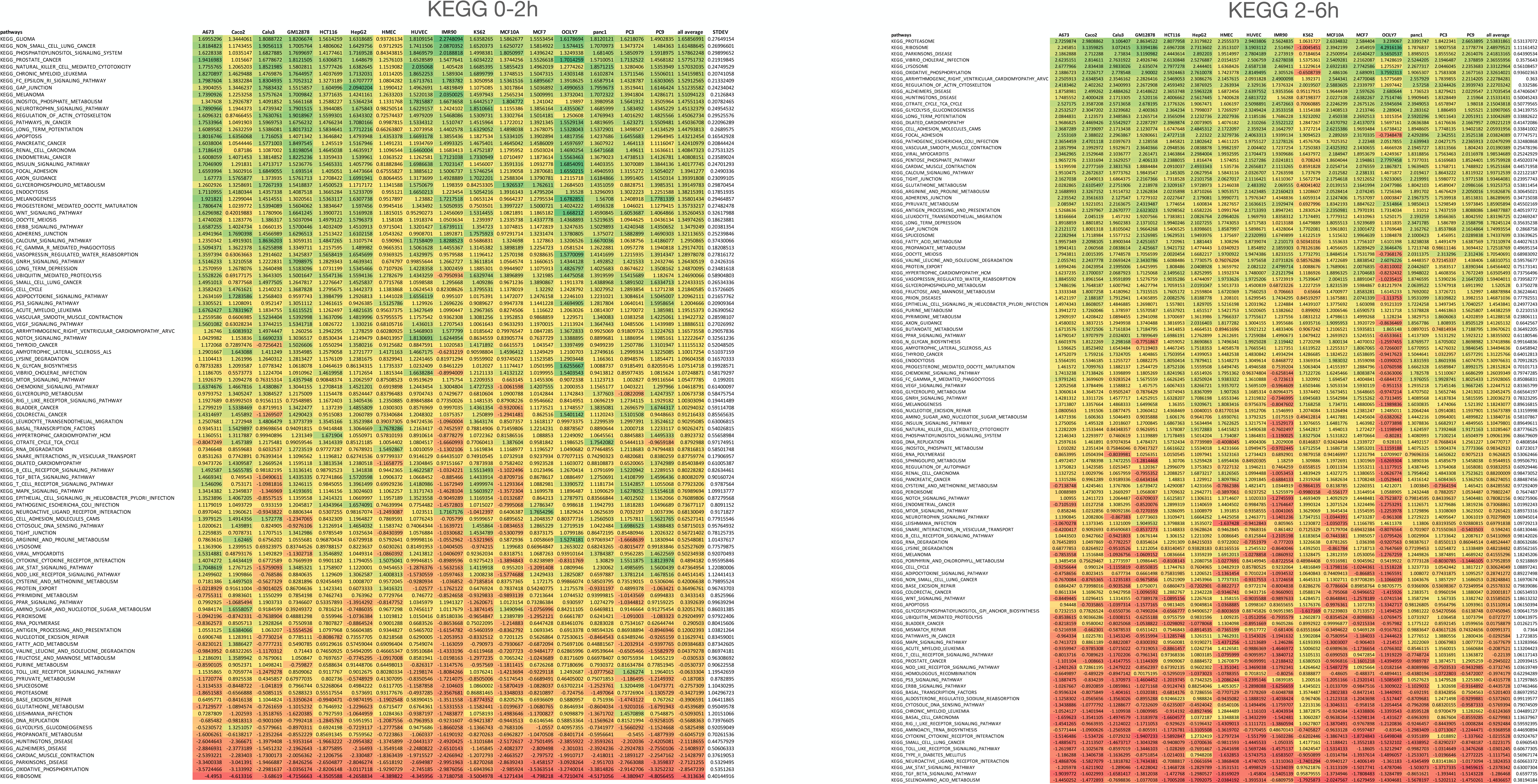
Co-regulation of RNA stability for transcripts with similar functions. GSEA analysis showing the Normalized Enhancement Score (NES) of relative stability of transcripts across 16 cell lines at 0-2h (left) and 2-6h (right). The different KEGG pathways (rows) were averaged for all 16 cell lines and this average was used to order the different pathways from the most positive (green) to the most negative (red). The standard deviation of averages of the NES values of the 16 cell lines are shown in the column on the right.

**Extended Figure 2c.**
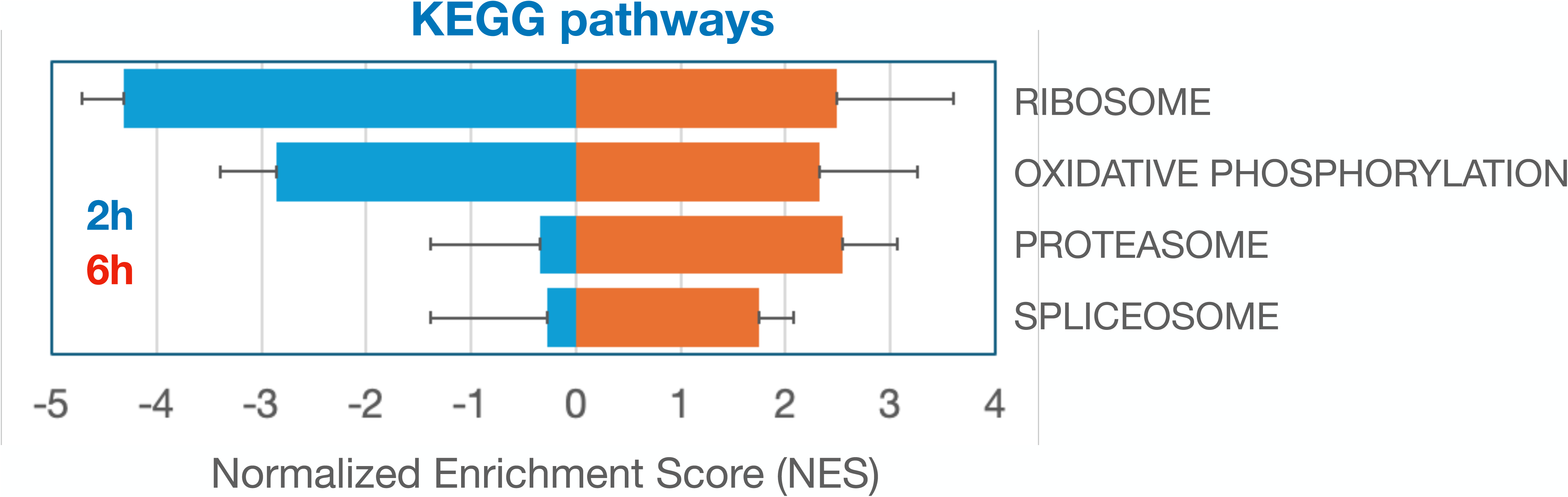
Normalized enrichment scores (NES) for relative transcript stability averaged across 16 cell lines for the KEGG pathways indicated. The error bars indicate the standard deviation of NES scores from the 16 cell lines. Related to Figure 2l.

**Extended Figure 3.**
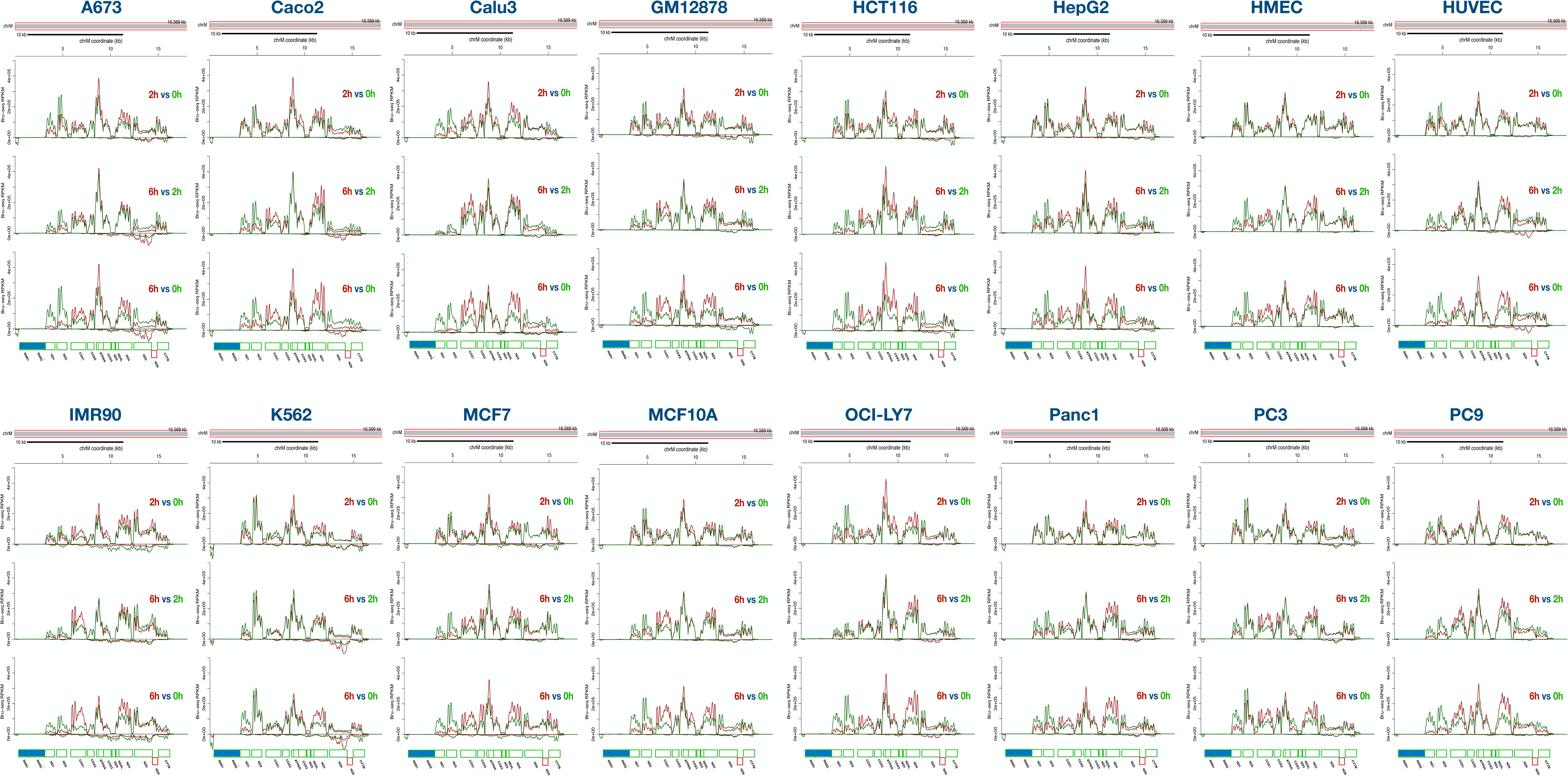
Mitochondrial transcripts show differential turnover rates. Read coverages across the mitochondrial genome are displayed for 2h vs 0h (top), 6h vs. 2h (middle) and 6h vs. 0h (bottom) across 16 cell lines.

**Extended Figure 4.**
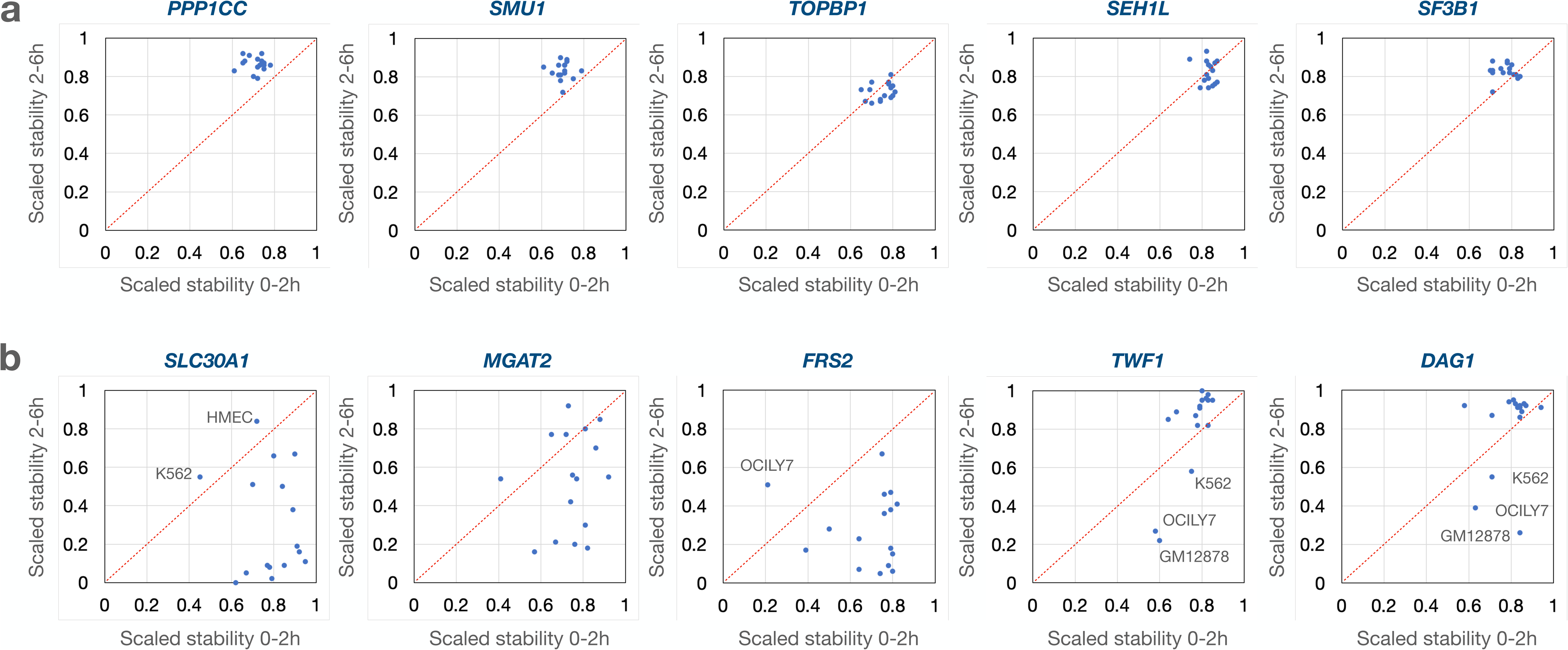
Transcript-specific vs. cell type-specific stability. **a**) Examples of transcripts that show very similar degradation patterns across all 16 cell lines. **b**) Examples of transcripts that show diverse degradation patterns across the 16 cell lines.

**Extended Figure 5.**
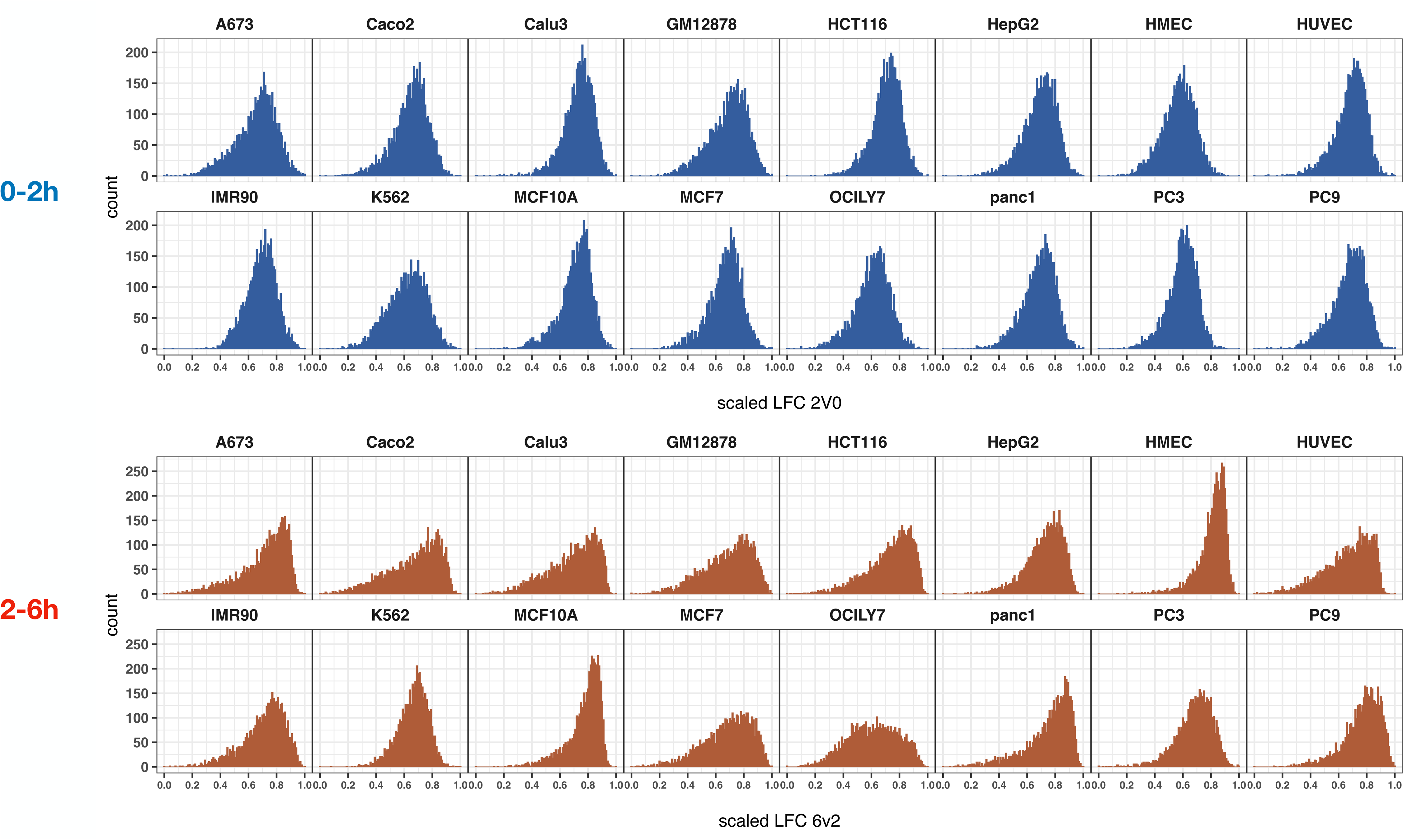
Distribution of Log2FC scaled stability values across the 16 cell lines for the 0-2h time period (top in blue) and for the 2-6h time period (bottom in red).

**Extended Figure 6.**
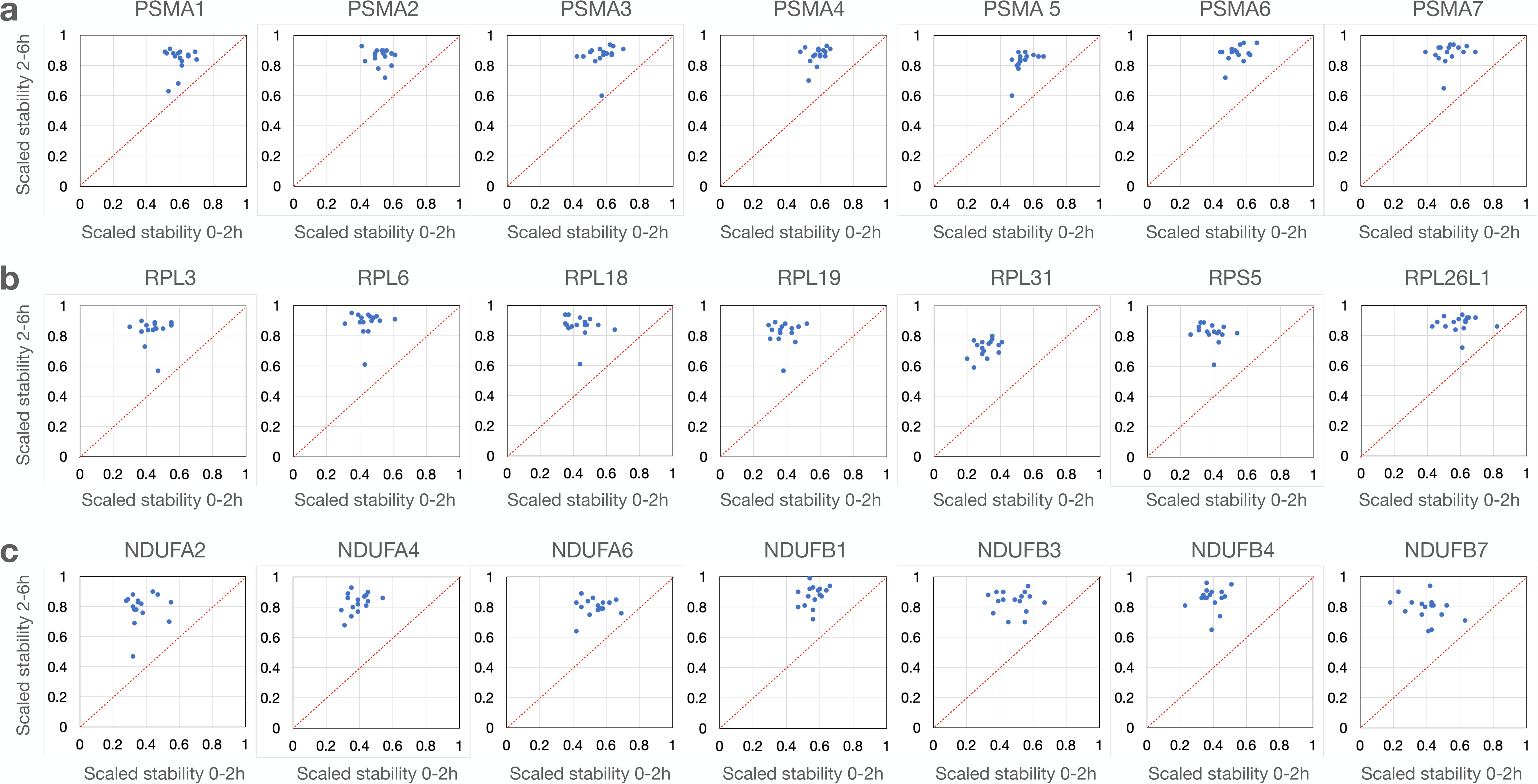
Initial high degradation rates followed by high stability for transcripts encoding proteins assembled into **a**) proteasomes **b)**, ribosomes and **c)** mitochondria.

**Extended Figure 7.**
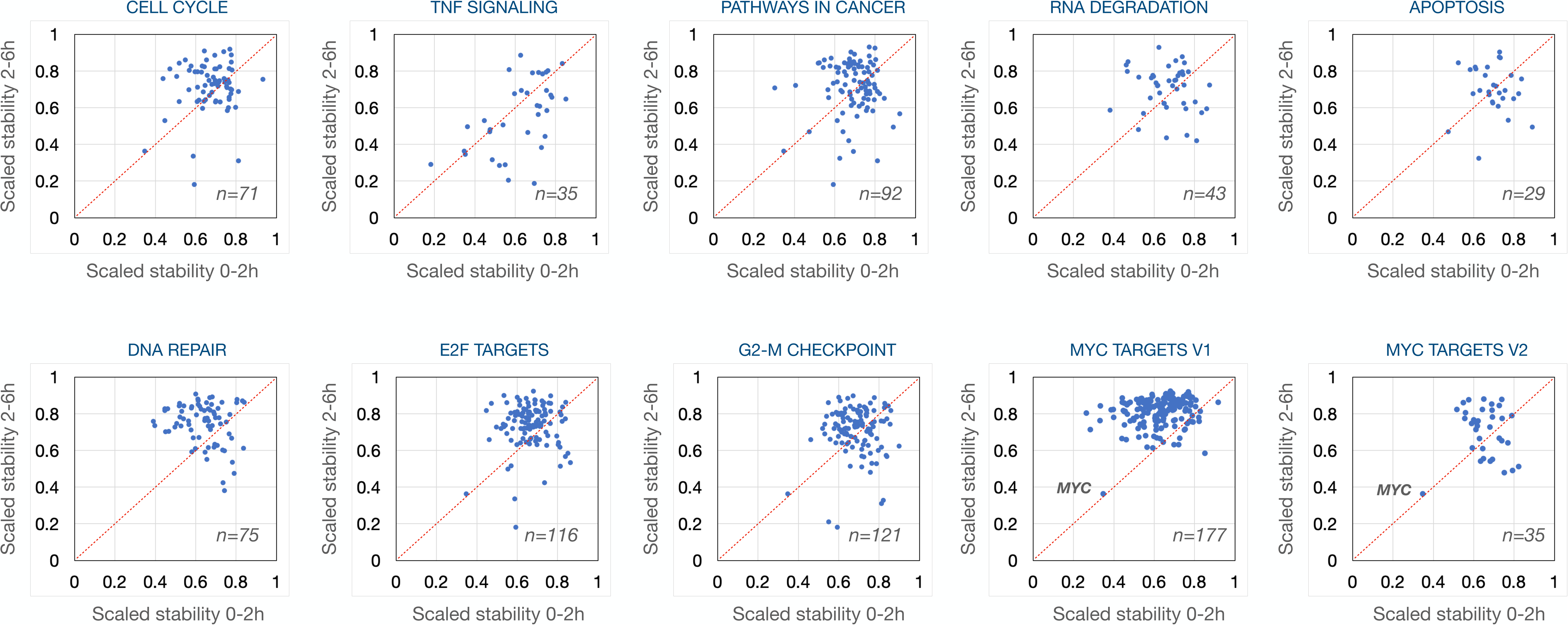
Distribution of scaled stability values for transcripts belonging to GSEA pathways. The scaled stability values of each individual transcript were averaged across the 16 cell lines and plotted in a 2-dimensional matrix with number of genes sampled in the pathway indicated.

**Extended Figure 8.**
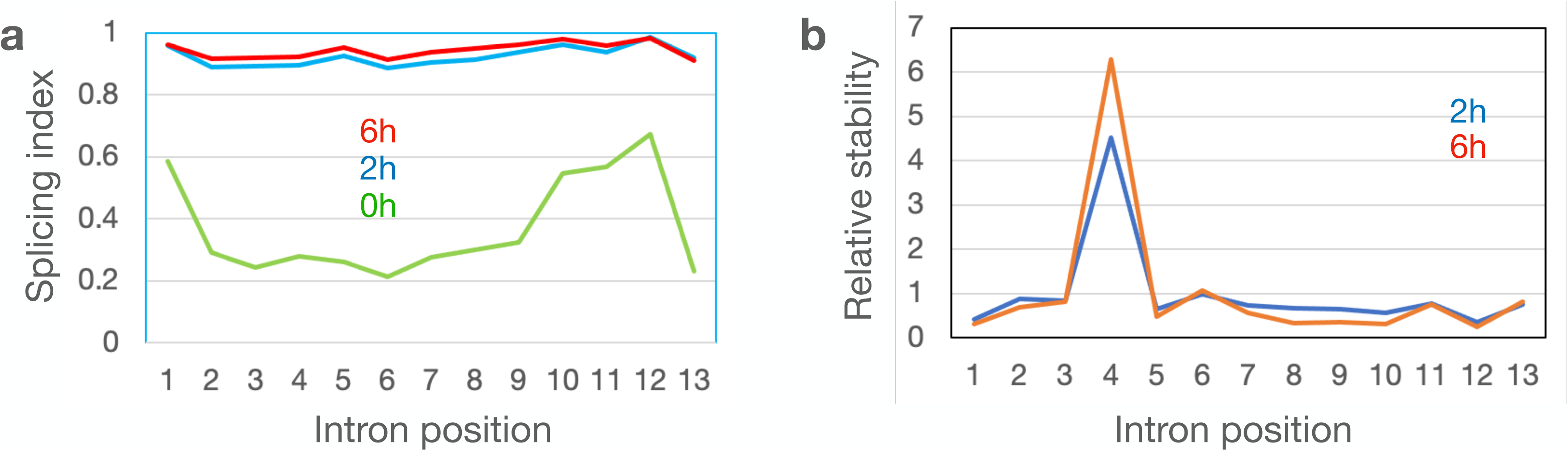
Retention of intron #4 in the *STAM* transcript despite efficient splicing across all 16 cell lines. (Related to Figure 5 c&d). **a)** The splicing indices across the *STAM* transcript at 0h, 2h and 6h were averaged from the 16 cell lines and plotted across the gene. **b)** The relative stabilities of the different introns across the *STAM* gene at 2h (0-2h) and 6h (2-6h) were averaged and plotted across the gene.

